## Supplementary Figures and Tables for "Mechanism of electroneutral sodium/proton antiporter from transition-path shooting"

**This file includes:**
**Extended Data Figure 1-13**
**Extended Data Table 1-3**

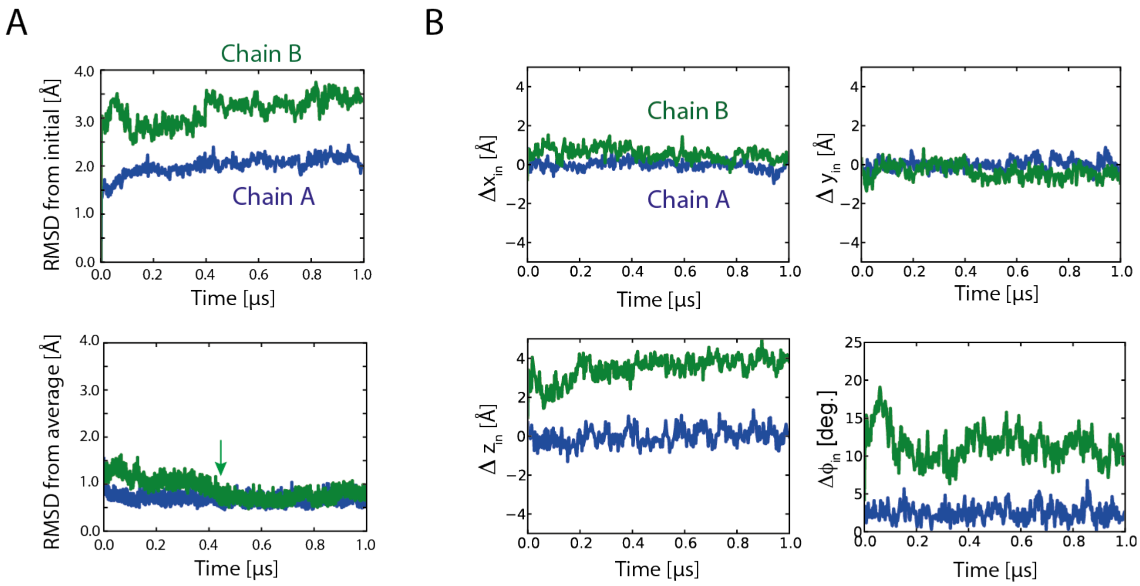


**Extended Data Figure 1 | Equilibrium MD simulation of asymmetric dimer following targeted MD simulation.** (A) RMSD of backbone atoms (Cα, C, O, N) from (top) the initial and (bottom) the average structures over the last 0.5 μs. The green arrow indicates the relaxation of protomer B towards the final structure at ~0.4 μs, associated with helix-5 motion. (B) Translational and rotational motions of the six-helix-bundle domain relative to the average inward-open structure.

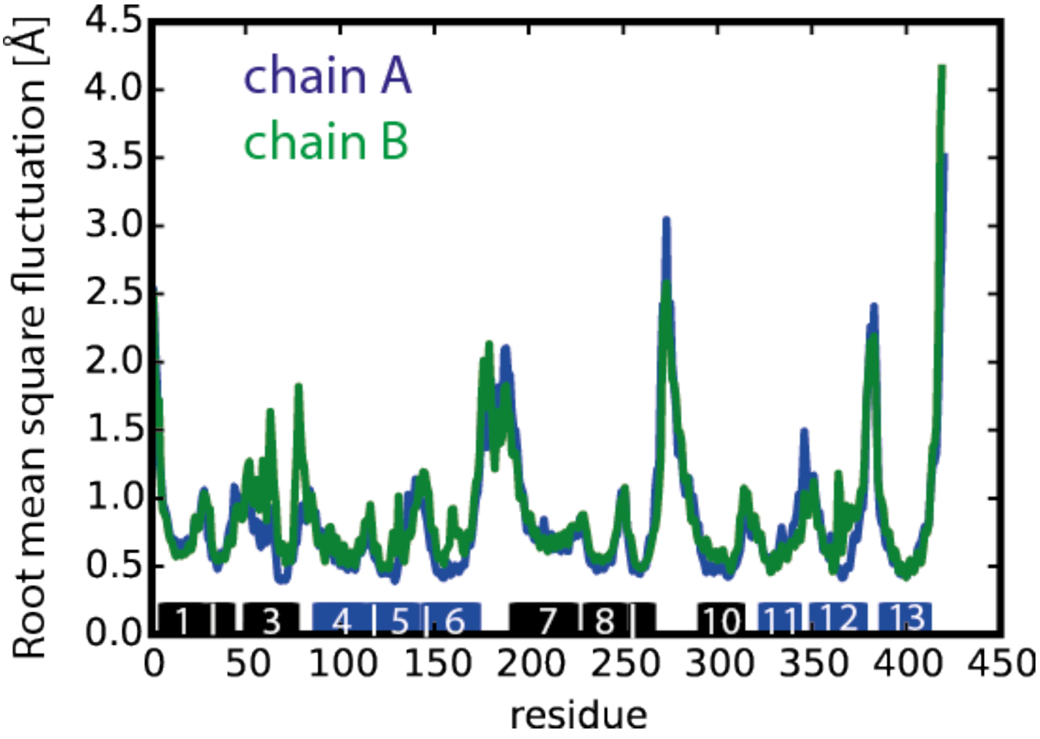


**Extended Data Figure 2 | Root-mean-square fluctuations of the Cα backbone in the asymmetric dimer.** The fluctuation profiles were obtained as a function of residue number during the second half of the free simulation of the asymmetric dimer. In the 1-$\mu$s simulation, the chain-A and chain-B protomers assumed inward- and outward-open conformations, respectively. The bars on the bottom show the helical regions used for structural superposition; numbers in the bars represent the helix number, coloured in blue and black for the six-helix-bundle and dimerization domains, respectively.

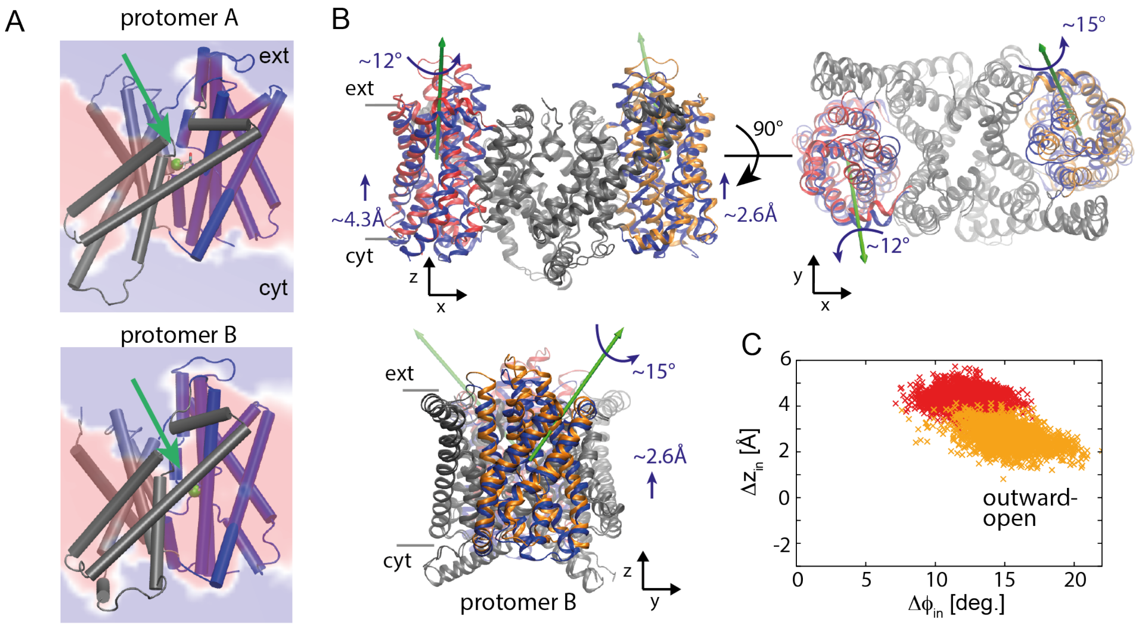


**Extended Data Figure 3 | Symmetric outward-open structure from targeted MD and 1-$\mu$s free simulation.** (A) Outward-open structures of protomers A and B with a section through the average water density in the 1-$\mu$s free MD simulation of the symmetric dimer (red: no water; blue: water at bulk density). The green sphere indicates H$\delta$ of the protonated Asp159. (B) Average dimer structure superimposed onto the average inward-open structure in (top) front and (bottom) side views. The green arrows show the axes of rotations of the six-helix-bundle domains. (C) Angle and z-coordinate changes of the six-helix-bundle domains of protomers A and B relative to the average inward-open structures (red and orange crosses, respectively).

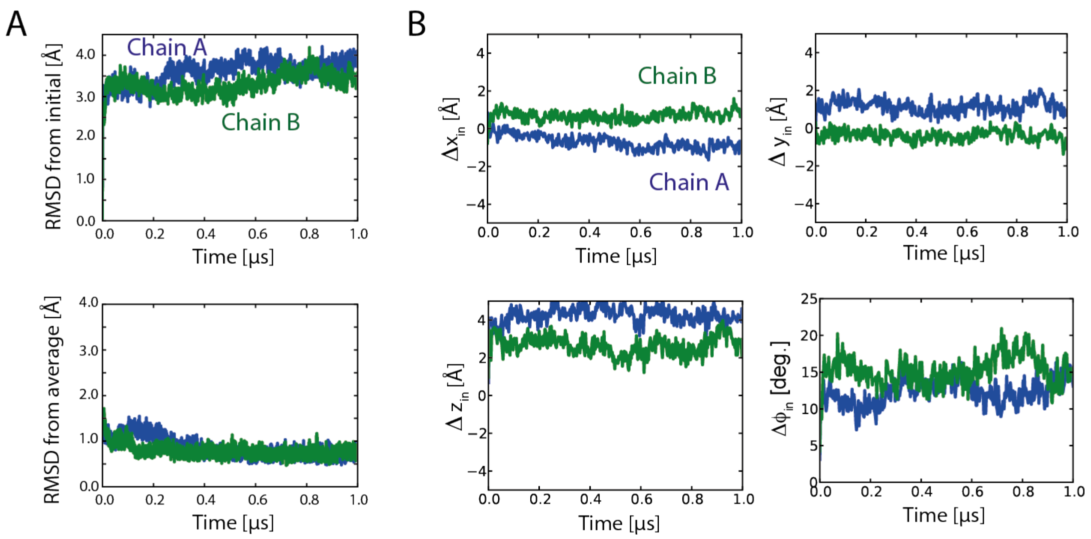


**Extended Data Figure 4 | Equilibrium MD simulation of symmetric dimer following targeted MD simulation.** (A) RMSD of backbone atoms (C $\alpha$ , C, O, N) from the initial (top) and average structures (bottom; averaged over the last 0.5 $\mu$ s). (B) Translational and rotational motions of the six-helix-bundle domain relative to the average inward-open structure.

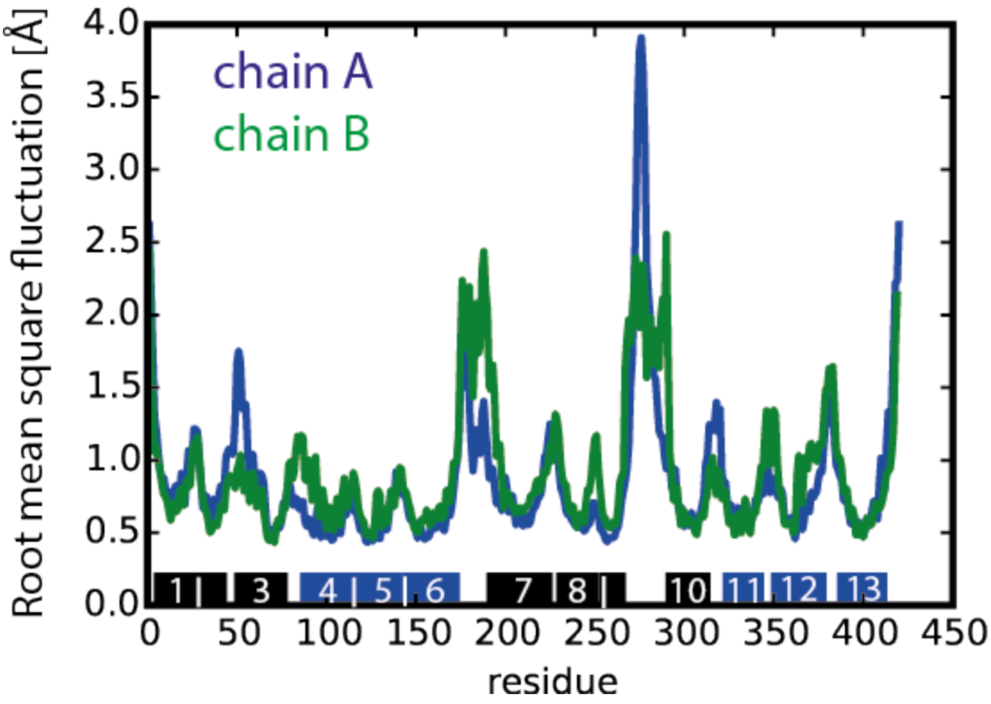


**Extended Data Figure 5 |** Root-mean-square fluctuations of the Cα backbone in the symmetric dimer. In the 1-μs simulation, both protomers assumed outward-open conformations. The bars on the bottom show the helical regions used for structural superposition; numbers in the bars represent the helix number, coloured in blue and black for the six-helix-bundle and dimerization domains, respectively.

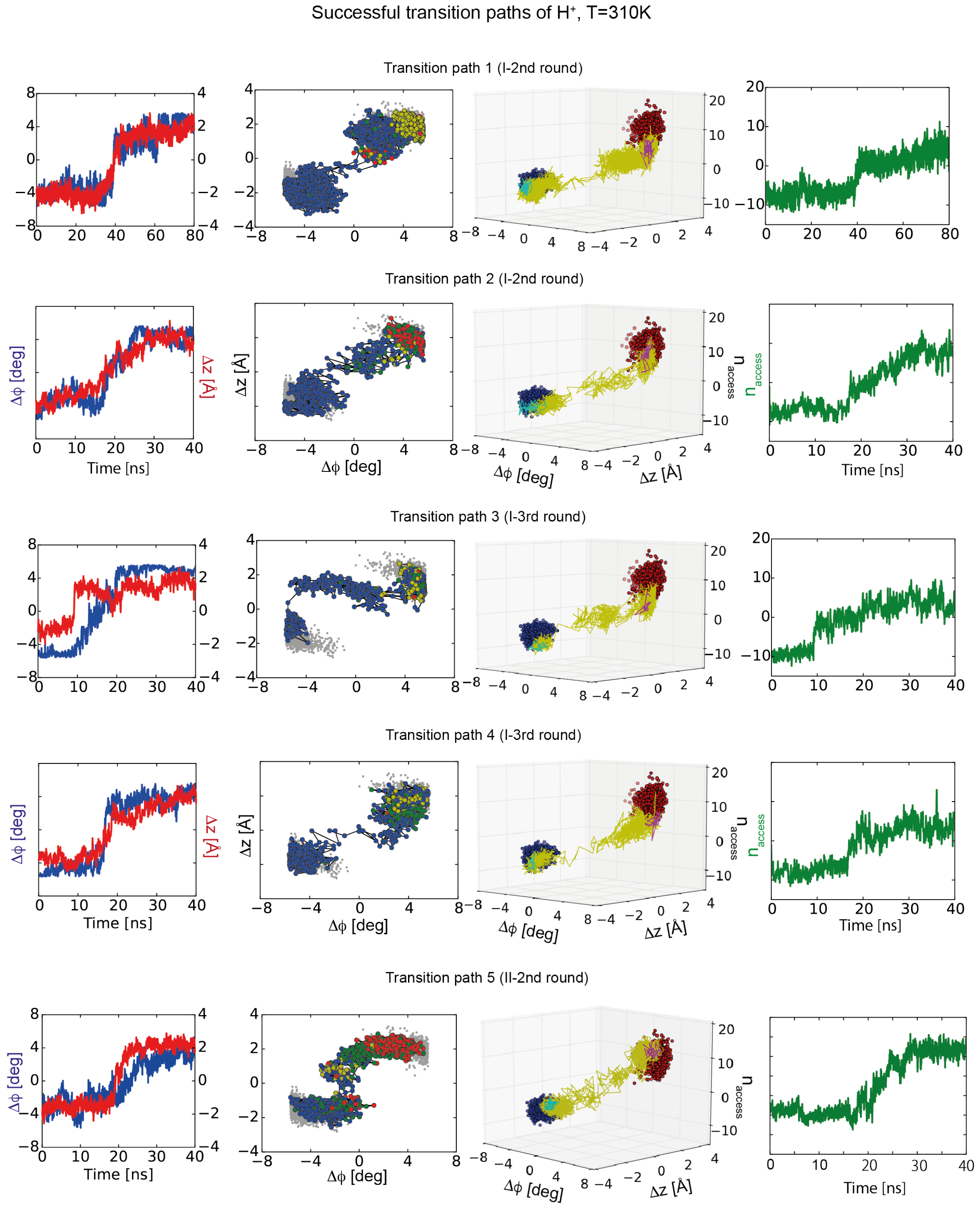


**Extended Data Figure 6 | Transition paths with protonated Asp159 at 310 K.**

Column 1 shows time series of the order parameters (blue/left axis: $\Delta\varphi$; red/right axis: $\Delta z$). Column 2 shows a projection of equilibrium runs (gray) and transition paths onto the $\Delta\varphi$ - $\Delta z$ plane. Inward-open, occluded, connected, and outward-open states are shown as blue, green, yellow, and red dots, respectively. In column 3, the hydration order parameter $n_{access}$ is included as a third coordinate, with the corresponding time series shown in column 4.

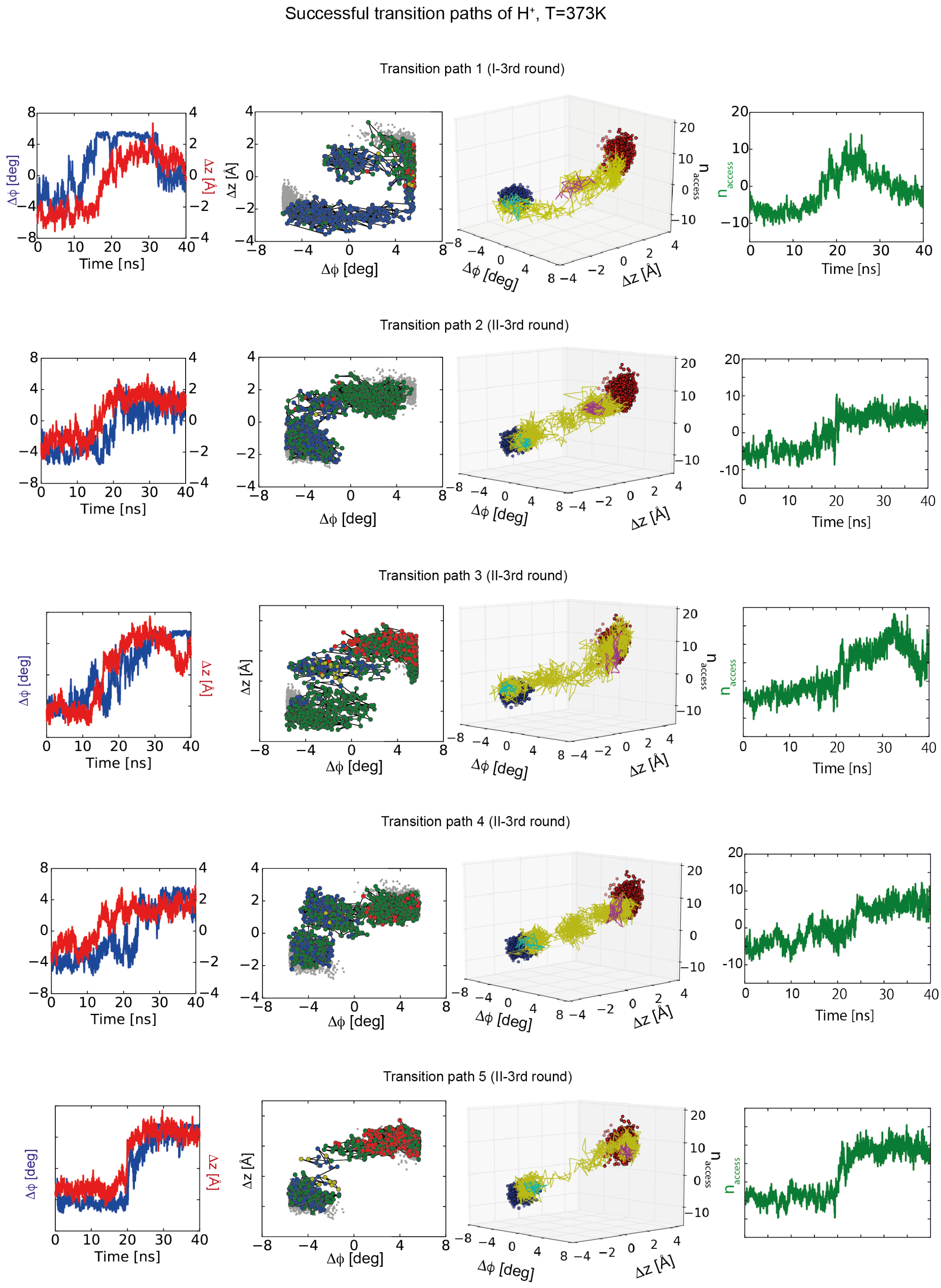


**Extended Data Figure 7 | Transition paths with protonated Asp159 at 373 K.**

Successful transition paths with protonated Asp159 at 373 K. Column 1 shows time series of the order parameters (blue/left axis: Δ$\varphi$; red/right axis: Δ$z$). Column 2 shows a projection of equilibrium runs (gray) and transition paths onto the Δ$\varphi$-Δ$z$ plane. Inward-open, occluded, connected, and outward-open states are shown as blue, green, yellow, and red points, respectively. In column 3, the hydration order parameter $n_{access}$ is included as a third coordinate, with the corresponding time series shown in column 4.

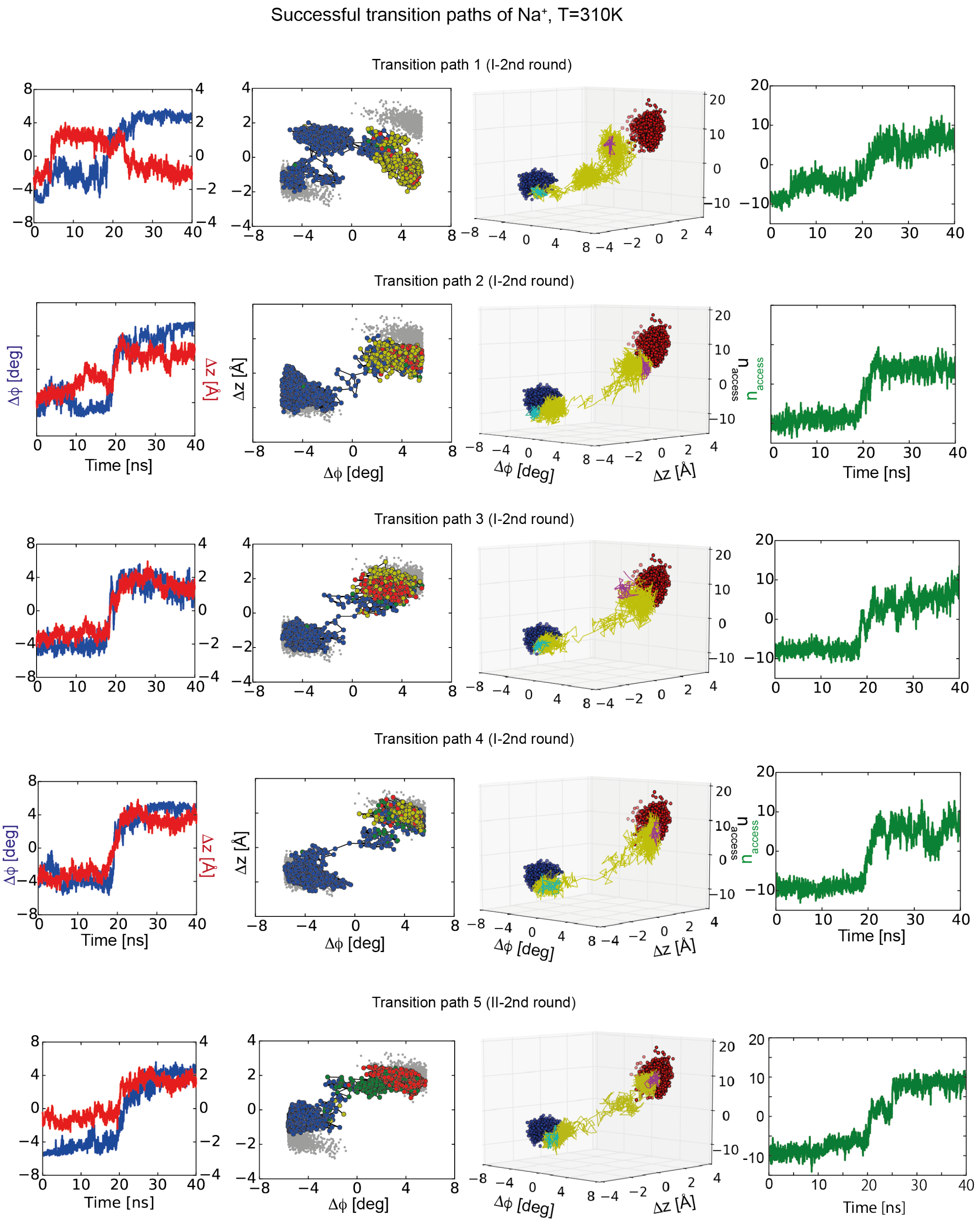


**Extended Data Figure 8 | Transition paths with $Na^+$ bound at 310 K.** Column 1 shows time series of the order parameters (blue/left axis: $\Delta\varphi$; red/right axis: $\Delta z$).

Column 2 shows a projection of equilibrium runs (gray) and transition paths onto the $\Delta\varphi$ - $\Delta z$ plane. Inward-open, occluded, connected, and outward-open states are shown as blue, green, yellow, and red points, respectively. In column 3, the hydration order parameter $n_{access}$ is included as a third coordinate, with the corresponding time series shown in column 4.

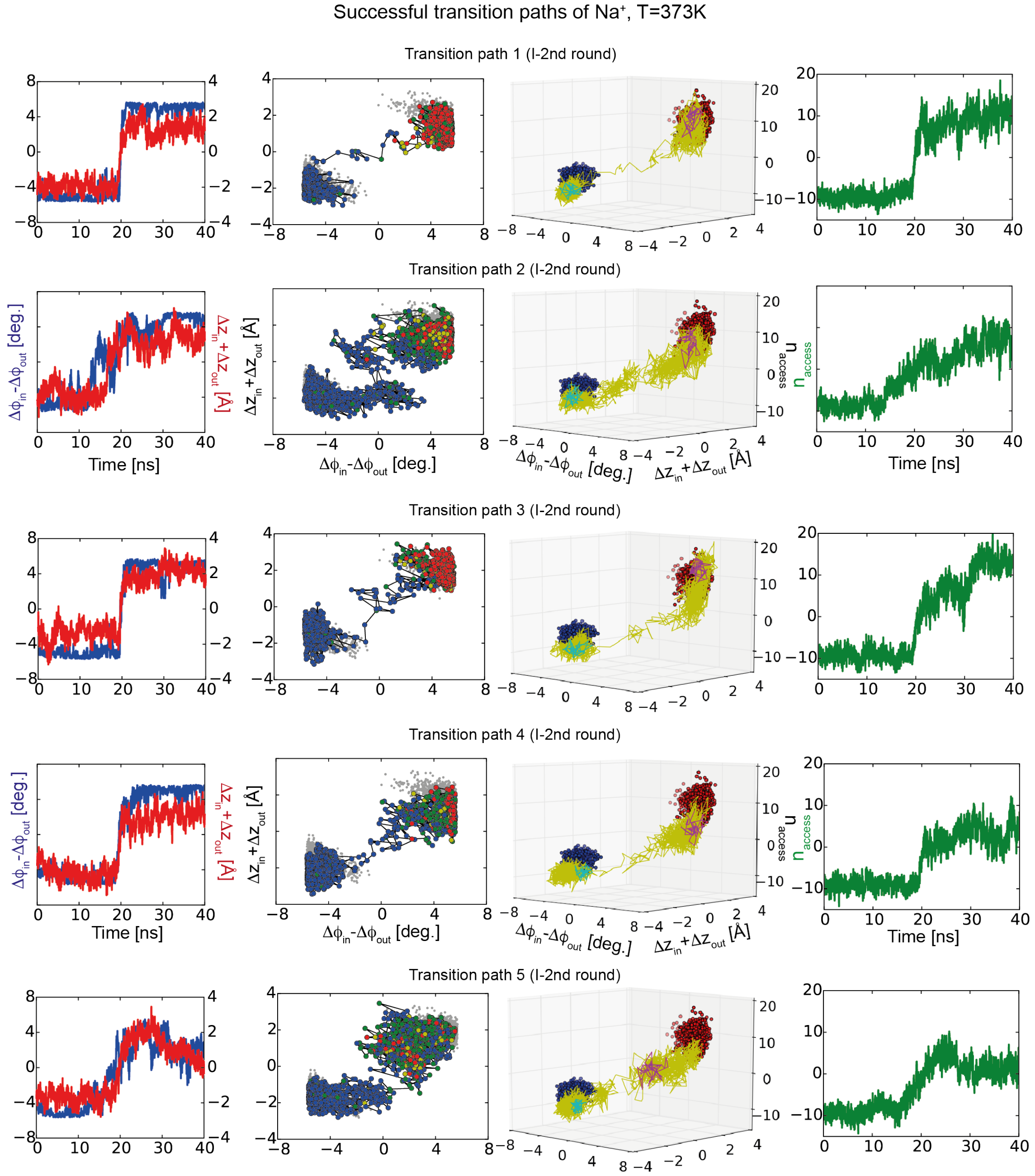


**Extended Data Figure 9 | Transition paths with $Na^+$ bound at 373 K.** Column 1 shows time series of the order parameters (blue/left axis: $\Delta\varphi$; red/right axis: $\Delta z$). Column 2 shows a projection of equilibrium runs (gray) and transition paths onto the $\Delta\varphi$ - $\Delta z$ plane. Inward-open, occluded, connected, and outward-open states are shown as blue, green, yellow, and red points, respectively. In column 3, the hydration order parameter $n_{access}$ is included as a third coordinate, with the corresponding time series shown in column 4.

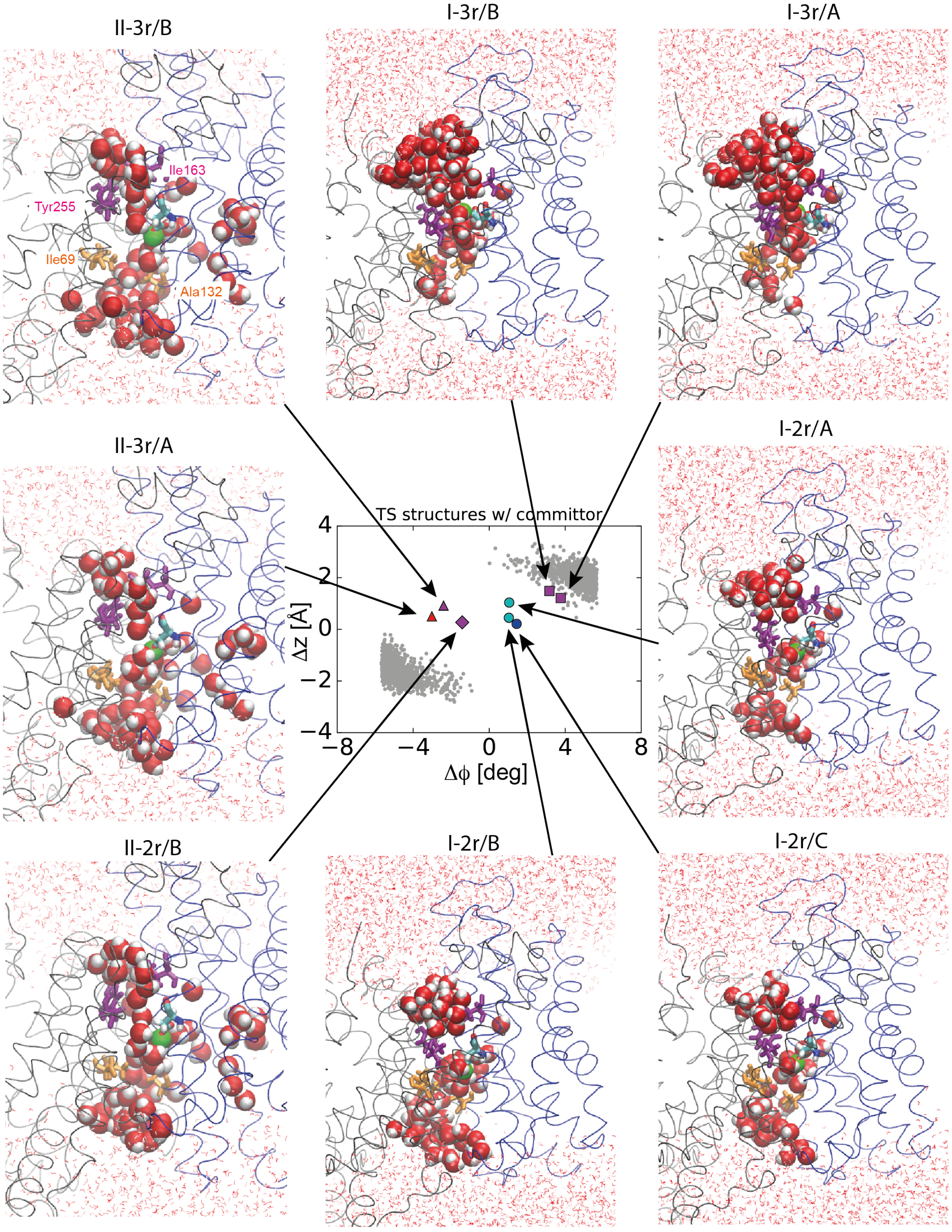


**Extended Data Figure 10 | Transition states.** Top and bottom rows: Gallery of eight transition states from rounds 2 to 3 of transition-path shooting starting from initial path I and II, respectively. Orange and magenta sticks represent the inward and outward gate residues, respectively. Asp159 is shown in stick representation and the bound $H^+$ as a green sphere. Water molecules within 15 Å of the bound $H^+$ of Asp159 are shown as spheres; other water molecules are shown as lines, and the protein backbone is shown as a tube. Centre: The transition-state structures are mapped onto the $\Delta\varphi$-$\Delta z$ surface. The structures for the second and third rounds of transition-path shooting starting from initial path I are shown as circles and squares, respectively. The structures for the second and third rounds of transition-path shooting following the initial path II are shown as diamonds and triangles, respectively. Estimated committor probabilities to the outward-open state are indicated by colour: >60% (red), 40-60% (magenta), 20-40% (cyan), <20% (blue). The gray dots represent $\Delta\varphi$-$\Delta z$ of structures sampled during last half of the 1 $\mu$s equilibrium simulation of the asymmetric dimer (protomer A: bottom left; protomer B: top right).

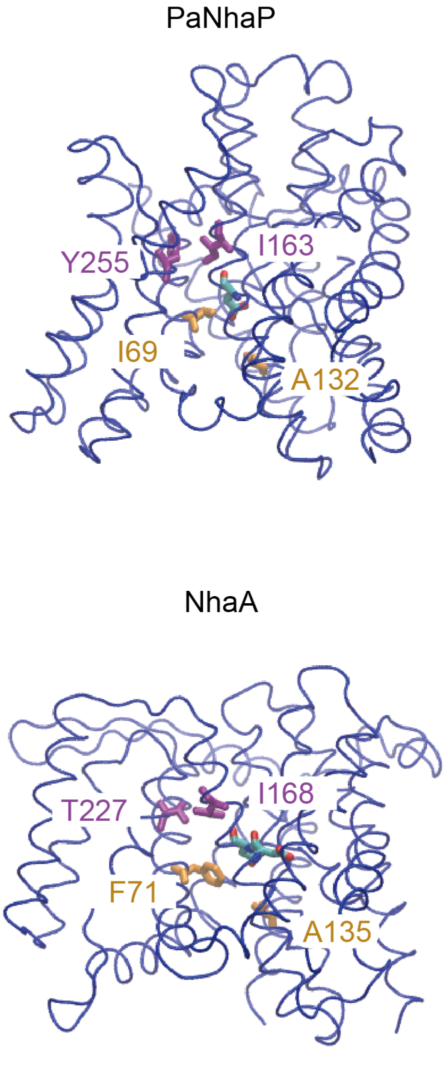


**Extended Data Figure 11 | Predicted gate residues from structural alignment.**

Outside and inside gate residues are shown in magenta and orange, respectively.

Conserved aspartates are coloured by atom.

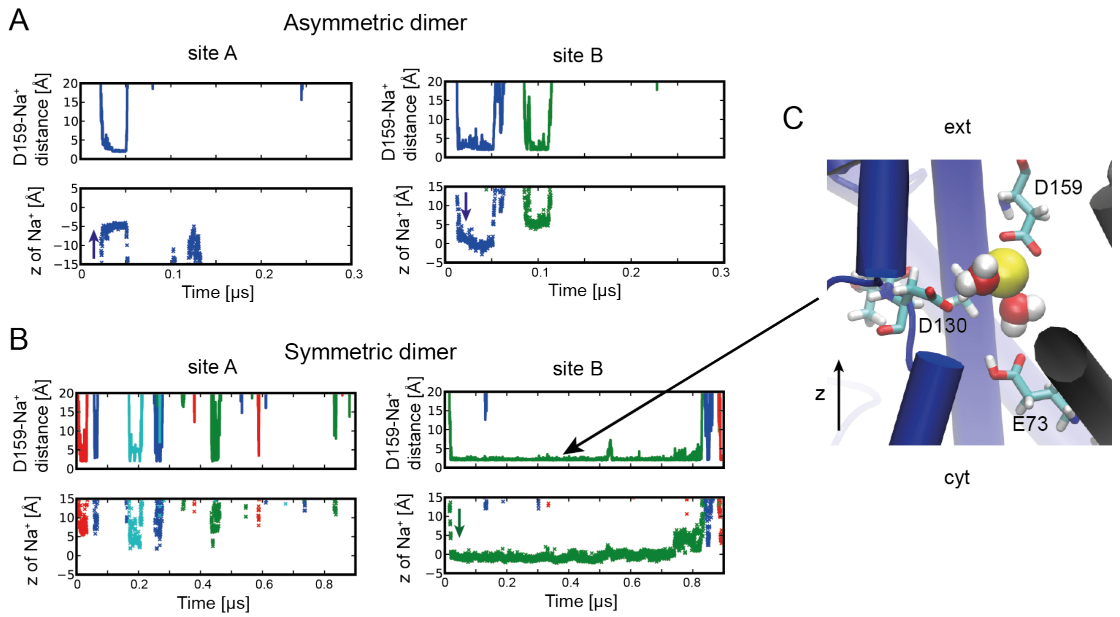


**Extended Data Figure 12 | Ion-binding events in simulations with Asp159 ionized.**

(**A**, **B**) Distances between $Na^+$ and O $\delta$ atoms of Asp159 (top), and z coordinates of the $Na^+$ (bottom) for site-A (left) and site-B (right). Trajectories of $Na^+$ ions are shown only when their distance from Asp159 is below 3 Å and remains bound for at least 4 ns. (**A**) Trajectories for the asymmetric dimer. (**B**) Trajectories for the symmetric dimer. (**C**) Snapshot of bound $Na^+$ (yellow sphere) in the outward-open state of the symmetric dimer trajectory.

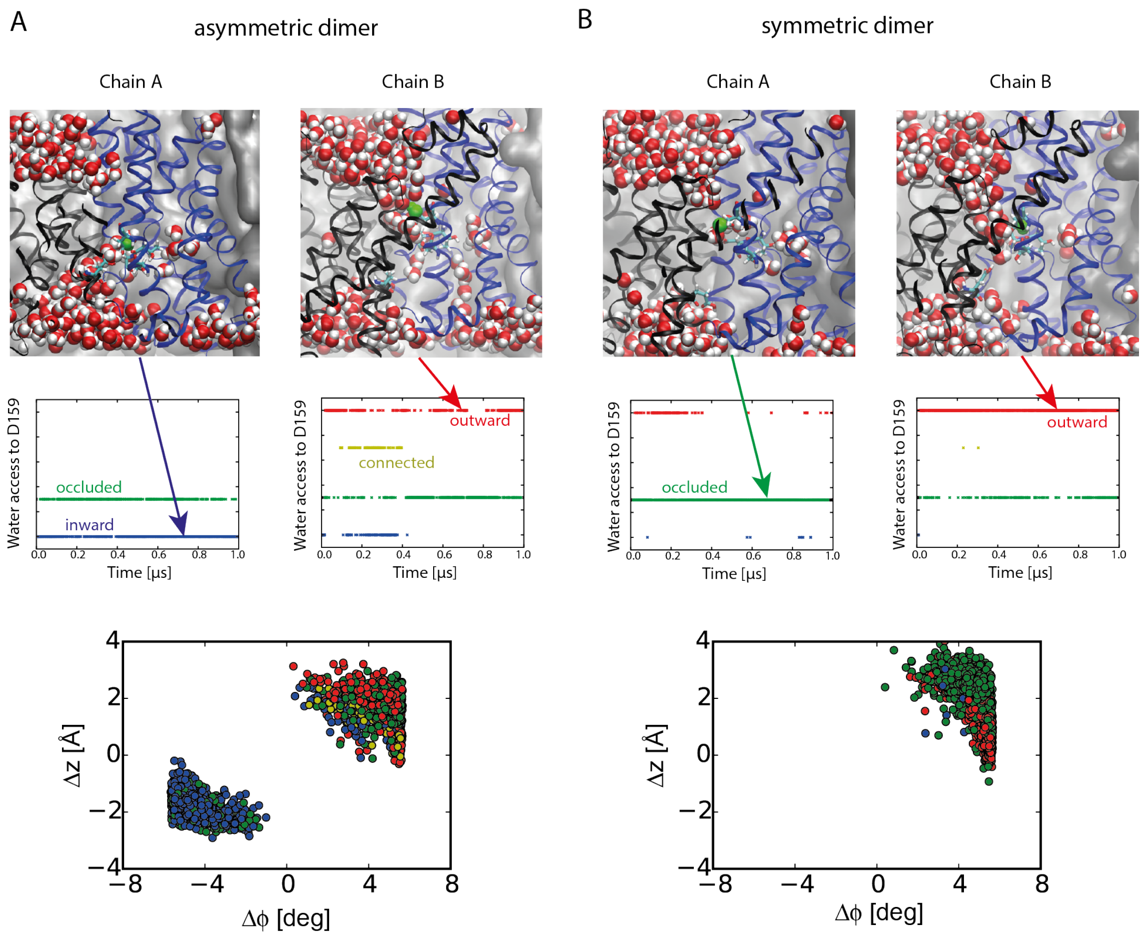


**Extended Data Figure 13 | Water accessibility to the ion-binding site.** Water molecules are clustered into groups by their pairwise distance, and the cluster of water molecules interacting with the carboxyl O$\delta$ atom of D159 (green) is analysed. (**A**) Asymmetric dimer and (**B**) symmetric dimer. (Top) Typical snapshots with protein backbone as ribbons, water molecules as red (oxygen) and white spheres (hydrogen), and lipids in surface representation. Middle row: Water access state during the 1-$\mu$s trajectories, with four distinct states: inward-open (blue), occluded (green), connected (gold), and outward-open (red). Bottom row: Projection of 1-$\mu$s trajectories onto $\Delta\varphi$-$\Delta z$ plane reporting on transporter domain motion. Symbol colours indicate water access state.

| | $H^+$, T=310K | $H^+$, T=373K | $Na^+$, T=310K | $Na^+$, T=373K | | |
|---|---|---|---|---|---|---|
| TS candidates | traj/in/out/tp | traj/in/out/tp | traj/in/out/tp | traj/in/out/tp | **$\Delta\phi$** | **$\Delta z$** |
| Initial path I | | | | | | |
| I-1r/6ns | 5/5/0/* | | | | 2.57 | 1.2 |
| I-1r/7ns | 5/3/0/* | | | | 2.73 | 1.24 |
| I-1r/8ns | 5/3/0/* | | | | 3.38 | 1.4 |
| I-1r/9ns | 5/1/2/* | | | | 3.03 | 1.7 |
| I-1r/10ns | 5/0/3/* | | | | 3.51 | 1.71 |
| I-2r/A | 36/22/6/1 | 14/13/0/0 | 12/8/2/2 | 12/4/7/2 | 1.05 | 1.04 |
| I-2r/B | 14/7/7/1 | 8/4/1/0 | 12/6/4/2 | 12/9/2/2 | 1.03 | 0.45 |
| I-2r/C | 8/4/0/0 | 8/4/1/0 | 12/8/0/0 | 12/9/1/1 | 1.44 | 0.21 |
| I-3r/A | 2/1/1/1 | | | | 3.77 | 1.21 |
| I-3r/B | 10/2/6/1 | 4/3/1/1 | | | 3.17 | 1.48 |
| I-4r/A | 4/4/0/0 | 8/4/2/0 | | | 3.05 | -0.1 |
| Initial path II | | | | | | |
| II-1r/6ns | 5/0/5/* | | | | 0.46 | 1.49 |
| II-1r/8ns | 5/0/5/* | | | | -2.17 | 1.29 |
| II-1r/9ns | 5/0/0/* | | | | -2.00 | 0.45 |
| II-1r/9.2ns | 5/1/0/* | | | | -2.52 | -0.17 |
| II-1r/9.3ns | 5/2/1/* | | | | -2.28 | -0.44 |
| II-1r/9.5ns | 5/3/1/* | | | | -2.62 | -0.24 |
| II-1r/10ns | 5/5/0/* | | | | -2.70 | -0.26 |
| II-2r/A | 10/0/3/0 | | 10/0/6/0 | | -1.73 | 0.92 |
| II-2r/B | 10/5/1/1 | 10/1/7/0 | 10/7/1/1 | | -1.42 | 0.26 |
| II-2r/C | 10/0/3/0 | | | | -1.57 | 1.39 |
| II-2r/D | 10/0/3/0 | | | | -0.08 | 1.58 |

| | | | | | | |
|---|---|---|---|---|---|---|
| II-2r/E | 10/0/4/0 | | | | 2.22 | 0.99 |
| II-3r/A | 10/2/2/0 | 10/2/4/1 | | | -3.02 | 0.50 |
| II-3r/B | 10/2/2/0 | 10/5/4/3 | | | -2.40 | 0.92 |
| II-3r/C | 10/6/2/0 | 10/0/3/0 | | | -1.98 | 0.29 |
| II-3r/D | 10/0/5/0 | 10/1/5/0 | | | -1.19 | 1.17 |
| II-3r/E | 10/0/4/0 | 10/0/4/0 | | | 0.58 | 1.17 |

**Extended Data Table 1. Transition-path shooting statistics.** Column 1 labels transition-state candidates, defined as configurations from which trajectories were initiated, according to the four/three rounds of the iterative transition-path shooting following initial paths I/II (I-1r to I-4r / II-1r to II-3r). The configurations of the first round are labelled in addition by the time point at which they were sampled in the TMD trajectory. In column 2-5, the number of total trajectory shots (traj), trajectories that end up in the inward-open state (in), outward-open state (out), and successful transition paths (tp) are listed, respectively, for states with $H^+$ and $Na^+$ bound, and for normal and elevated temperatures. The star indicates that in the first round, trajectory segments were not initiated with sign-inverted initial velocities and could thus not be stitched together to form true transition paths. In column 6 and 7, the rotational angles (degrees) and *z*-translations (Å) of the six-helix-bundle domain are listed.

| | H$^+$, T=310K | | H$^+$, T=373K | | Na$^+$, T=310K | | Na$^+$, T=373K | |
|---|---|---|---|---|---|---|---|---|
| | **# traj** | **duration** | **# traj** | **duration** | **# traj** | **duration** | **# traj** | **duration** |
| eq in | 1 | 1.0 | 1 | 0.4 | 1 | 0.4 | 1 | 0.3 |
| eq a-out | 1 | 1.0 | | | 1 | 0.5 | | |
| eq s-out | 1 | 1.0 | | | 1 | 1.0 | | |
| Initial path I | | | | | | | | |
| 1r TPS | 25 | 0.01 | | | | | | |
| 2r TPS | 92/22 | 0.04/0.02 | 30 | 0.02 | 36 | 0.02 | 36 | 0.02 |
| 3r TPS | 24 | 0.02 | 4 | 0.02 | 24 | 0.02 | 10 | 0.02 |
| 4r TPS | 16 | 0.02 | 8 | 0.02 | | | | |
| Initial path II | | | | | | | | |
| 1r TPS | 35 | 0.02 | | | | | | |
| 2r TPS | 50 | 0.02 | 10 | 0.02 | 20 | 0.02 | | |
| 3r TPS | 50 | 0.02 | 50 | 0.02 | | | | |
| Total | 10.87 μs | | 2.44 μs | | 3.5 μs | | 1.22 μs | |

**Extended Data Table 2. Computational effort.** The equilibrium and transition-path shooting trajectories are listed as "eq" and "*n*r TPS", respectively, where *n* represents *n*-th round of transition-path shootings. "in", "a-out," and "s-out" refer to inward-open, asymmetric inward/outward-open, and symmetric outward-open states, respectively. All times are in units of microseconds. The total simulation time was 18 μs.

| Name | PDB | Conserved ASP | Outside gate | Inside gate |
|---|---|---|---|---|
| **PaNhaP** | 4CZA | D159 | I163, Y255 | I69, A132 |
| | | | V365, L369 | L70, P131 |
| **MjNhaP1** | 4CZB | D161 | I165, Y248 | I69, A134 |
| | | | V350, L354 | L70, P133 |
| **NhaA** | 1ZCD | D163, D164 | I168, T227 | F71, A135 |
| | | | F344, I345 | F72, I134 |
| **NapA** | 5BZ2 | D156, D157 | L161,I238,V239 | L67, G129 |
| | | | I337 | L68, V128 |

**Extended Data Table 3. Gate residues predicted from structural alignment of inward-open conformations.** For each structure, the main gate residues are in the upper line, while complementing residues are in the lower line.
